# Association of diet and inflammation with the vaginal microbiota of pregnant individuals with or without IBD

**DOI:** 10.1101/2024.04.23.590846

**Authors:** Daniela Vargas-Robles, Yan Rou Yap, Biplab Singha, Joyce Tien, Mallika Purandare, Mayra Rojas-Correa, Camilla Madziar, Mellissa Picker, Tina Dumont, Heidi Leftwich, Christine F. Frisard, Doyle V. Ward, Inga Peter, Barbara Olendzki, Ana Maldonado-Contreras

**Affiliations:** Department of Microbiology and Physiological Systems, Program of Microbiome Dynamics; School of Medicine; University of Massachusetts Chan Medical School, Worcester, MA, 01655; Department of Genetics and Genomic Sciences, Icahn School of Medicine at Mount Sinai, New York, NY, 10029; Department of Obstetrics and Gynecology, Division of Maternal-Fetal Medicine, University of Massachusetts Chan Medical School, Worcester, MA, 01655; Department of Population and Quantitative Health Science. University of Massachusetts Chan Medical School, Worcester, MA, 01655

**Keywords:** Vaginal, microbiota, IBD, inflammation, pregnancy

## Abstract

**Background and aims:** Vaginal dysbiosis has been associated with adverse pregnancy outcomes. Here, we characterized the vaginal microbiota of pregnant individuals with inflammatory bowel disease (IBD) and investigated whether gut or vaginal inflammation and diet influence the vaginal microbiota diversity of these individuals.

**Study Design:** We recruited 48 individuals in their third trimester of pregnancy (IBD=23 and HC=18). We characterized the vaginal microbiota by *16S rRNA* sequencing and the gut microbiota by shotgun sequencing. We measured fecal calprotectin in stool and pro-inflammatory cytokines in vaginal fluids. We determine dietary quality using validated 24-hour dietary recalls.

**Results:** Pregnant individuals with IBD exhibit higher levels of fecal calprotectin and increased expression of Th17 pro-inflammatory cytokines (i.e., IL-6, IL-8, IL-17) in the vaginal mucosa compared to healthy pregnant individuals. High fecal calprotectin correlated with high vaginal microbiota diversity. Also, IL-4 (reduced in IBD) was associated with vaginal microbial composition. Regardless of IBD status, pregnant individuals with healthier diets and particularly optimal servings of vegetables and sugars exhibited a vaginal microbiota dominated by *Lactobacillus crispatus*, a species associated with a lower risk of preterm birth and bacterial vaginosis.

**Conclusion:** Besides gut inflammation, pregnant individuals with IBD also exhibit a Th17 immune tone in the vaginal mucosa. The vaginal microbiota diversity or composition, particularly high in the beneficial *L. crispatus,* is positively associated with healthier diets, regardless of IBD status.

**Why was the study conducted?:** An altered vaginal microbiota has been implicated in preterm birth. There is no research on the vaginal microbiome and the factors that influence it in pregnant individuals with Inflammatory Bowel Disease (IBD) at a higher risk of preterm delivery.

**Key findings:** Pregnant individuals with IBD exhibit a comparable vaginal microbiome to healthy pregnant individuals. However, pregnant individuals with IBD present a vaginal immune profile characterized by increased levels of Th17 pro-inflammatory cytokines. High dietary quality, and optimal consumption of vegetables and added sugars were associated with vaginal dominance by the beneficial *L. crispatus*.

**What does this add to what is known?:** Our results indicate that the vaginal immune environment and not the microbiome might explain poor pregnancy outcomes for individuals with IBD. Moreover, our study supports the importance of diet to favor *L. crispatus,* a bacterium associated with a lower risk of preterm birth.

## Introduction

The vaginal microbiota composition of pregnant individuals with IBD is stable throughout pregnancy but has not been directly compared to a healthy cohort (1). Only one study has reported that pregnant individuals with IBD were more likely to have vaginal infections compared to healthy pregnant individuals (2). In general, healthy pregnancies are characterized by vaginal microbiotas with low bacterial diversity and dominant *Lactobacillus* species (3). For instance, pregnant individuals with a vaginal microbiota dominated by *L. crispatus* have a lower risk of preterm birth compared to individuals with low-lactobacilli abundance (4). Moreover, vaginal microbial communities dominated by *L. crispatus* are better protected against infections than those dominated by *L. iners* (5). Therefore, understanding *Lactobacillus* species dominance in individuals with IBD is important to predicting their risk of poor pregnancy outcomes. Our goal is to characterize and compare the vaginal microbiota of pregnant individuals with and without IBD using high-throughput microbiota sequencing.

Moreover, we sought to determine the role of environmental factors, such as diet and local inflammation on the vaginal microbiota composition. To our knowledge, there have been only a few studies evaluating the influence of diet on the vaginal microbiota of pregnant individuals using high throughput microbiota sequencing (6–8), yet none of the studies included all the relevant diet components from validated instruments aiming at measuring dietary intake. Additionally, the immune tone of the vaginal mucosa in pregnant women with IBD remains understudied. Hence, we will test whether diet influences the vaginal microbiota makeup thus diet can be potentially used as a strategy to revert vaginal dysbiosis during pregnancy, and whether an inflammatory environment in the gut and vagina are linked to vaginal microbiota profiles.

## Materials and Methods

### Recruitment

We conducted a case–control study nested into our ongoing MELODY Trial (9) including participants with 27^th^–29^th^ weeks of gestation before any dietary intervention. Pregnant women with and without IBD were recruited nationwide under approved IRB protocol (IRB # H00016462) as previously described (10) (see supplementary methods). IBD disease activity was evaluated using validated scoring systems, namely the Harvey Bradshaw index (11) for participants with CD and the 6-point Mayo score (12) for participants with UC.

### Sample collection

Vaginal and stool samples were self-collected using the OMNIgeneVAGINAL collection tube (DNA Genotek, Canada) or the ALPCO EasySampler kit (ALPCO, USA) following manufacturer instructions. Samples were received in the lab frozen 30h after sample collection.

### Nucleic acid isolation

DNA from both vaginal and stool samples was isolated with Dneasy PowerSoil Pro kit (QIAGEN, Germany) and the RNA from vaginal samples was isolated with PowerMicrobiome kit (QIAGEN, Germany) following the manufacturer’s protocol.

### Cytokine expression

RNA from vaginal samples was subjected to qRT–PCRs (see supplementary methods). Oligonucleotides used to estimate cytokine expression are listed in **Table S1** (Integrated DNA Technology, USA).

**Fecal calprotectin quantification** was performed using the CalproLab ELISA ALP (Svar Life Sciences, Norway) according to the manufacturer’s instructions. Total protein was quantified using the Pierce BCA Protein Assay kit (Thermo Fisher Scientific, USA). Calprotectin was normalized to initial stool weight (ng calprotectin/mg stool).

**Vaginal microbiota sequencing and profiling** was performed by *16S rRNA* sequencing of the V3-V4 hypervariable region as previously described (13). Sequencing libraries were sequenced on 600 cycles using the MiSeq platform (Illumina, CA, USA). QIIME2 was used to process paired-end sequences. The DADA2 (14) algorithm was used for quality control and to obtain representative sequences (Amplicon Sequence Variant or ASV). We used a custom database (15, 16) for taxonomy classification. Only taxa with at least 0.10% abundance and present in a minimum of one sample were used for the analyses, as previously done (17, 18). **Table S2** describes the sequence counts included in the analyses.

**Gut microbiota sequencing and profiling** were performed using whole genome sequencing. Specifically, library generation and 150bp paired-end sequencing were carried out on the Illumina NextSeq 500 platform. KneadData (dec_v0.1, http://huttenhower.sph.harvard.edu/kneaddata) was used to eliminate human sequences and for quality control. MetaPhlAn4 database (vOct22)(19) was used for taxonomic assignment. **Table S2** describes the sequence counts included in the analyses.

**Vaginal and gut microbiota diversity analyses** were done in R, particularly the Phyloseq package (20). We performed data imputation for two individuals lacking BMI information (IBD=2), and four individuals (IBD=2, HC=2) without fecal calprotectin measurements using the median values for the IBD or HC group, respectively. All analyses were performed at the ASV level. Microbial alpha diversity was estimated using the Shannon and Simpson’s Indexes (1-D) with rarefied sequencing data. Shannon and Simpson’s Index were log-transformed to reach ‘normality’ of the residuals when necessary. To determine associations in alpha diversity we used linear regression models corrected by cofounders as age, body mass index (BMI), antibiotic use, and gestational diabetes. Results for the best-fitted model are reported.

We used Permutational Multivariate Analysis of Variance (PERMANOVA (21)) with adonis2 function to evaluate beta diversity (measured by Aitchison distance) of non-rarefied sequencing data.

We only assessed the associations between the vaginal microbiota and cytokines that were significantly different between IBD and HC. When assessing the association with dietary components, we only included dietary components with no collinearity (Spearman correlation >|0.5|). One participant did not complete dietary recalls and thus was excluded from diet analyses.

### Discriminant taxa analysis

Microbial taxa and their association with clinical variables were assessed using MaAsLin2’s (22), also including the previously named confounding variables.

### Community State Types (CST)

Each vaginal sample was classified into CST as described before (23). To compare CST by discrete variables (health status) we use Fisher exact test (24); and by continuous variables (i.e., fecal calprotectin, cytokine expression, or diet score) Kruskal-Wallis test.

### Dietary assessment

We conducted 24-hour dietary recalls (24HDRs) around the same time as vaginal/stool collection and estimated the Healthy Eating index-2015 (HEI-2015) as previously described by us (10) (see supplementary methods). Wilcoxon test was used to compare HEI-2015 and its dietary components by health status and CST.

## Results

A total of 48 pregnant individuals were enrolled in the study: 23 with diagnosed IBD (n=18 CD, and n=5, UC) and 25 HC without IBD. Participants’ demographics and clinical information are summarized in **Table 1**. Briefly, participants’ mean age was 33.8 years, most had normal BMI (41.7%) or were overweight (37.5%), and most self-identified as White (93.8%). Only a few participants reported gestational diabetes or the use of antibiotics currently or previously during pregnancy. None of the clinical and demographic variables differed by health status (IBD vs. HC, **Table 1**) or IBD diagnosis (CD vs. UC, **Table S3**). More than 50% of the IBD participants were in self-reported remission (**Table 1**).

**Table 1.** Demographic and clinical variables for pregnant individuals with Inflammatory Bowel Disease (IBD) or Healthy Controls (HC) recruited for the study between 2019 and 2022.

| Demographics and clinical variables | IBD<br>(N=23) | HC<br>(N=25) | Overall<br>(N=48) | P<br>value<sup>&</sup> |
| --- | --- | --- | --- | --- |
| **Age** |  |  |  | 0.414 |
| Mean (SD) | 33.3 (4.63) | 34.4 (4.97) | 33.8 (4.79) |  |
| Median [Min, Max] | 33.0 [22.0, 41.0] | 36.0 [22.0, 42.0] | 34.0 [22.0, 42.0] |  |
| **BMI categories <sup>&&</sup>** |  |  |  | 0.179 |
| Underweight | 1 (4.3%) | 0 (0%) | 1 (2.1%) |  |
| Normal | 12 (52.2%) | 8 (32.0%) | 20 (41.7%) |  |
| Overweight | 8 (34.8%) | 10 (40.0%) | 18 (37.5%) |  |
| Obese | 2 (8.7%) | 7 (28.0%) | 9 (18.8%) |  |
| **Race** |  |  |  | 0.490 |
| White | 23 (100%) | 22 (88.0%) | 45 (93.8%) |  |
| Asian | 0 (0%) | 1 (4.0%) | 1 (2.1%) |  |
| Other | 0 (0%) | 2 (8.0%) | 2 (4.2%) |  |
| **Gestational diabetes** |  |  |  | 0.098 |
| Yes | 3 (13.0%) | 0 (0%) | 3 (6.3%) |  |
| No | 16 (69.6%) | 21 (84.0%) | 37 (77.1%) |  |
| Information unavailable | 4 (17.4%) | 4 (16.0%) | 8 (16.7%) |  |
| **Use of antibiotic during all pregnancy** |  |  |  | 0.468 |
| No | 19 (82.6%) | 22 (88.0%) | 41 (85.4%) |  |
| Yes | 3 (13.0%) | 3 (12.0%) | 6 (12.5%) |  |
| Information unavailable | 1 (4.3%) | 0 (0%) | 1 (2.1%) |  |
| **IBD diagnosis** |  |  |  | - |
| Crohn's disease | 18 (78.3%) | 0 (0%) | 18 (37.5%) |  |
| Ulcerative colitis | 5 (21.7%) | 0 (0%) | 5 (10.4%) |  |
| Healthy controls | 0 (0%) | 25 (100%) | 25 (52.1%) |  |
| **IBD disease activity <sup>&&&</sup>** |  |  |  | - |
| Mild disease | 6 (26.1%) | 0 (0%) | 6 (12.5%) |  |
| Remission | 13 (56.5%) | 0 (0%) | 13 (27.1%) |  |
| Information unavailable | 4 (17.4%) | 25 (100%) | 29 (60.4%) |  |
| **Use of IBD medication** |  |  |  | 1.70E-06 |
| No | 9 (39.1%) | 25 (100%) | 34 (70.8%) |  |
| Yes | 14 (60.9%) | 0 (0%) | 14 (29.2%) |  |
| **Preterm** |  |  |  | 0.106 |
| No | 15 (65.2%) | 17 (68.0%) | 32 (66.7%) |  |
| Yes | 4 (17.4%) | 0 (0%) | 4 (8.3%) |  |
| Information unavailable | 4 (17.4%) | 8 (32.0%) | 12 (25.0%) |  |
| **Infant birth weight (g)** |  |  |  | 0.151 |
| Mean (SD) | 3150 (470) | 3330 (662) | 3240 (579) |  |
| Median [Min, Max] | 3230 [1810, 3710] | 3290 [1080, 4520] | 3230 [1080, 4520] |  |

<sup>&</sup>Fisher's exact test for categorical variables and Wilcoxon test for continuous variables.
<sup>&&</sup>BMI's categories correspond to the WHO's classifications: Underweight (<18.5), normal weight (18.5–24.9), overweight (≥25.0), and obese (≥30).
<sup>&&&</sup>Disease activity was estimated using the Harvey Bradshaw Index and the Mayo score for individuals with Crohn's Disease or Ulcerative colitis, respectively.

### Vaginal microbiota diversity is associated with gut and vaginal inflammation, regardless of health status

We first determined whether the vaginal microbiota diversity and composition differed by health status and found no significant differences (**Figure S1**). We also found no significant differences in the vaginal microbiota by IBD medications or when comparing individuals with IBD not on medication with the HC (**Figure S2**).

Given that most of the study participants were in remission or with mild disease we sought to further determine inflammatory markers that could influence the vaginal microbiota. We observed significantly higher levels of fecal calprotectin in IBD participants compared to their HC counterparts. Concomitantly, fecal calprotectin levels were positively associated with vaginal microbiota Simpson diversity but not with Shannon index (**Figure 1 and S1**) or for the beta diversity (**Figure S1**). Additionally, there were no significant associations between fecal calprotectin and vaginal microbiota diversity by health status (**Figure S1**).

**Figure 1.**
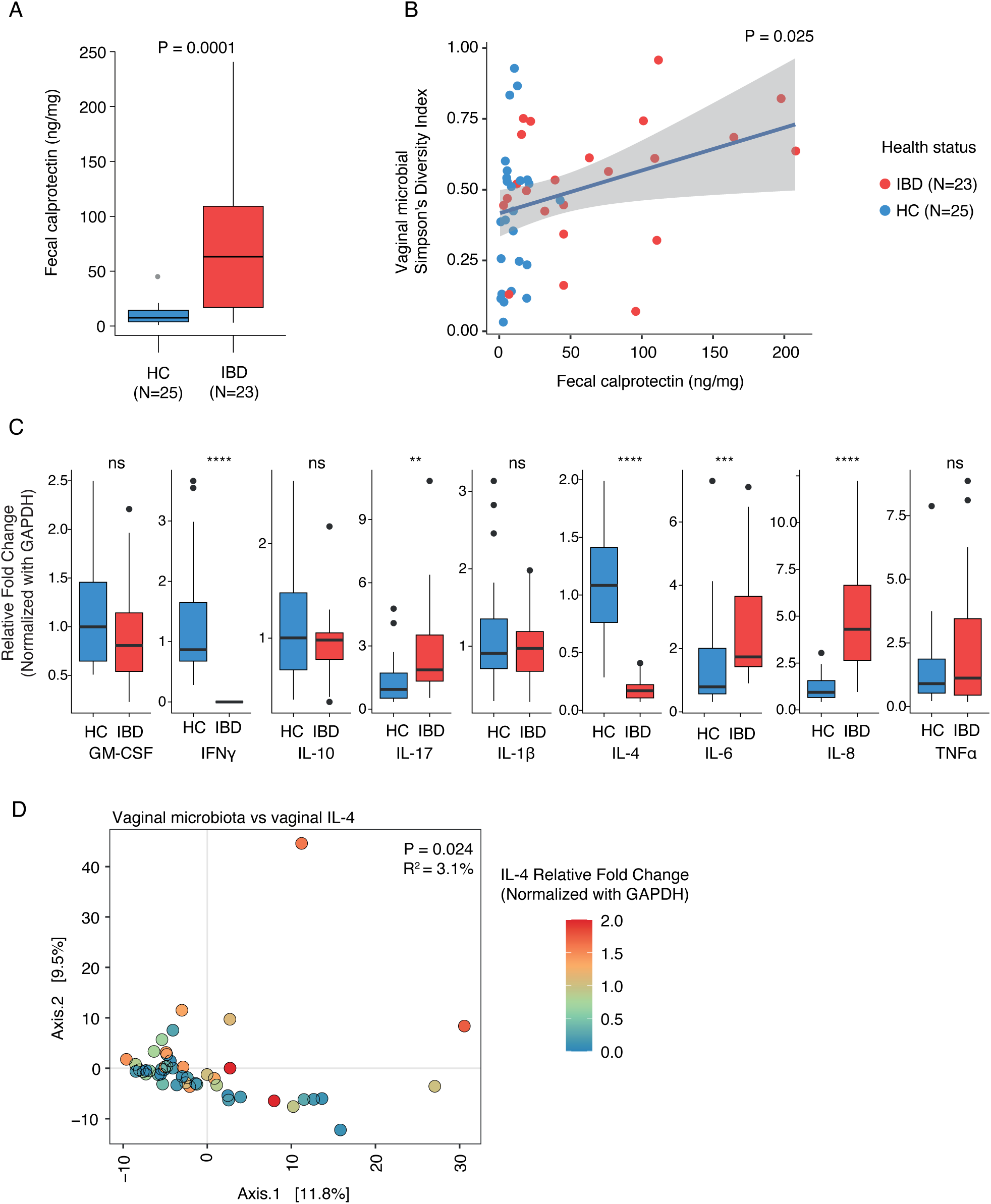
Variation in inflammatory markers and microbial diversity by health status. (A) Fecal calprotectin levels of individuals with Inflammatory Bowel Disease (IBD) or Healthy Controls (HC; Wilcoxon test). (B) Linear regression correlation between vaginal microbial alpha diversity and fecal calprotectin levels. (C) Expression of cytokines on the vaginal mucosa of IBD and HC participants (Wilcoxon test. Asterisks denote ** <5E-2, *** <5E-3, ****<5E-4). (D) Principal Coordinates Analysis (PCoA) of vaginal microbial beta diversity by IL-4 levels (Aitchison distance and PERMANOVA).

We then assessed inflammation in the vaginal mucosae. We observed that pregnant individuals with IBD exhibited higher gene expression of Th17 pro-inflammatory cytokines, specifically IL-6, IL-8, and IL-17 than HC. Conversely, expression of Th1 and Th2 cytokines IFN-γ and IL-4 respectively, was lower in pregnant individuals with IBD compared to HC (**Figure 1**). We observe that the vaginal microbiota composition – not its alpha diversity-correlated with the expression of IL-4 in the vaginal mucosa (**Figures 1 and S3**).

There were no specific bacteria associated with any of the inflammatory markers studied. We only identify *Dialister* sp. and WAL 1855D (an uncultured bacterium from the *Tissierellaceae* family) increased in pregnant individuals with high BMI (**Figure S4**).

Finally, we examined the gut microbiota of the pregnant individuals included in the study. There were no significant differences in the gut microbiota diversity and composition by health status or fecal calprotectin levels (**Figure S5**). However, we observed that *Collinsella* was lower in pregnant individuals with IBD compared to HC (**Figure S5**).

### Vaginal microbial diversity is associated with the consumption of vegetables and added sugars

Pregnant individuals in this cohort had a HEI-2015 score of 63.8, comparable to the average of 63.0 reported by pregnant individuals in the USA (25, 26). There were no significant differences in the HEI-2015 score by health status (**Table S5**). Similarly, there were no significant differences in vaginal microbial diversity or composition by HEI-2015 score (**Figure S6**).

We further investigated the associations between each dietary component included in the HEI-2015 score and the vaginal microbiota diversity. Consumption of each dietary component was similar between pregnant individuals with IBD and HC (**Table S5**). Whole fruit, fatty acid, and seafood/plant protein were excluded from the analysis due to collinearity (Spearman score >|0.5|). We observed that high scores of total vegetables (high vegetable intake) were predictive of a high vaginal microbiota diversity (**Figure 2**) along with increasing abundance of *L. crispatus* (**Figure 3**). *L. iners* showed an opposite trend, although not significant (**Figures 3 and S7**). We observed that vaginal microbial composition differed by added sugar score, although no bacterial taxon was significantly associated with it (**Figure 2**).

**Figure 2.**
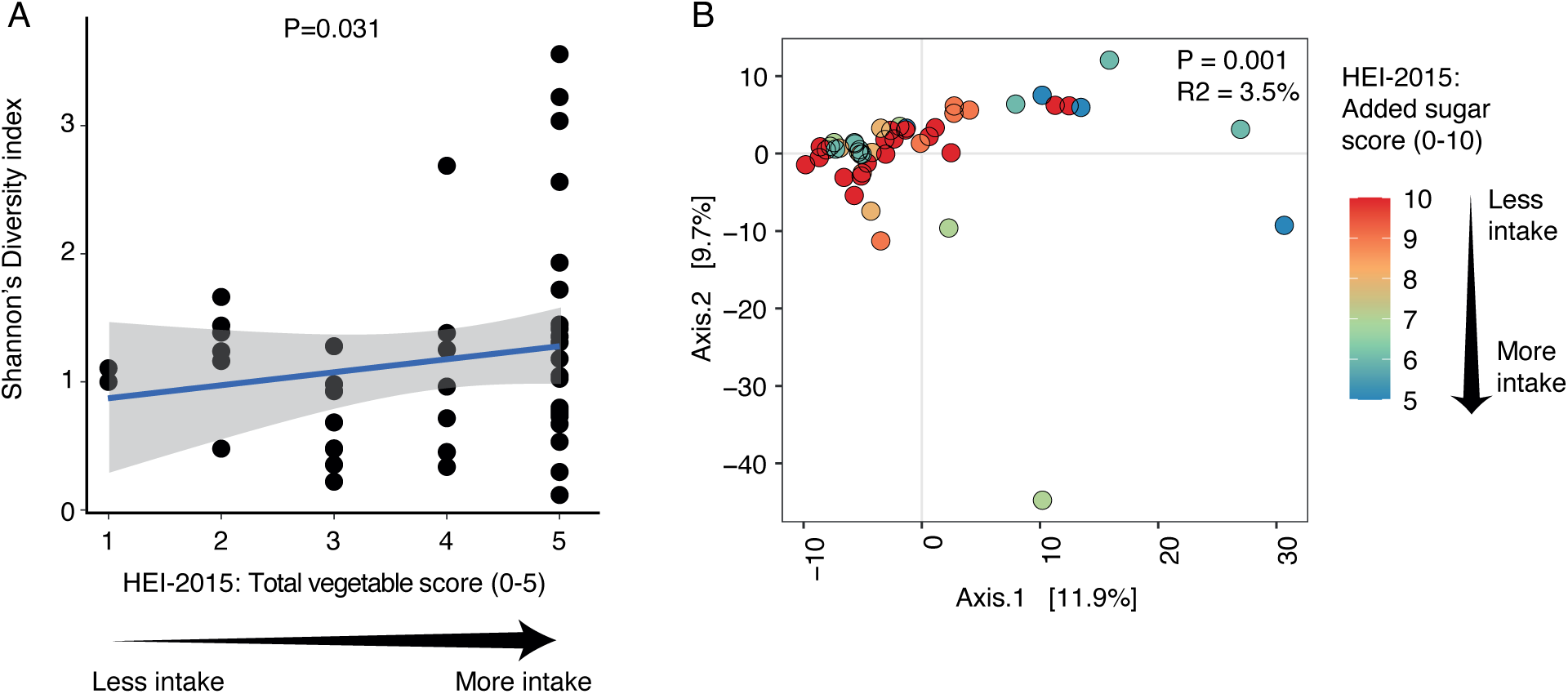
HEI-2015 total vegetable and added sugar components are significantly associated with vaginal microbiota diversity or composition, respectively. (A) Linear regression of microbial alpha diversity and HEI-2015 total vegetable score. Gray shades in the graphs represent the 95% confidence interval. (B) Principal Coordinates Analysis (PCoA) demonstrates significance for vaginal microbial composition by HEI-2015 added sugar score (Aitchison distance and PERMANOVA).

**Figure 3.**
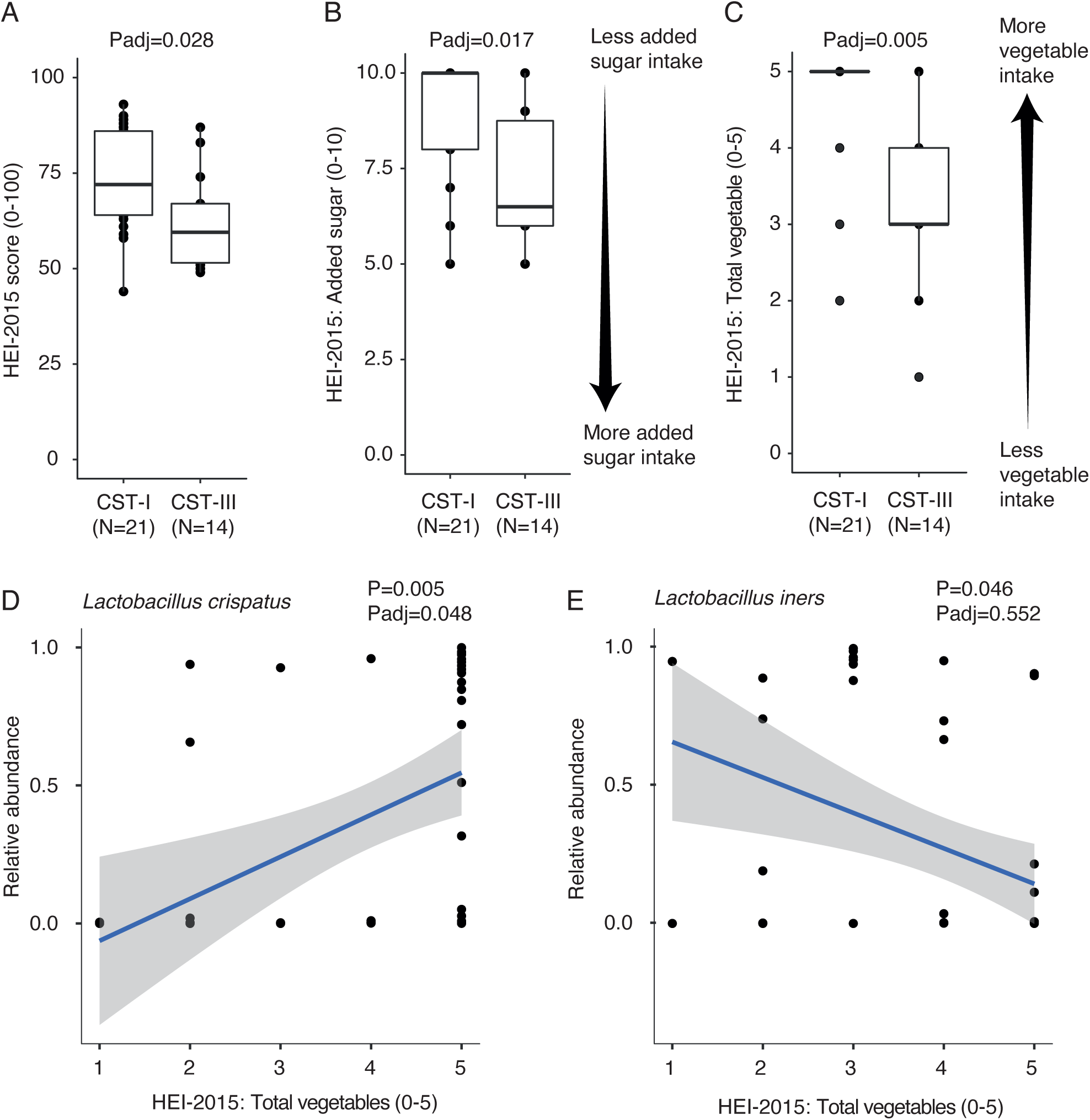
HEI-20215 added sugar and total vegetable correlate with CST classification. Pregnant individuals with CST-I (*L. crispatus*-dominated profile) exhibit higher scores for (A) HEI-2015, (B) added sugar, and (C) total vegetables (Wilcoxon-test, adjusting for multiple comparisons). The arrows indicate the direction of consumption of each dietary component. (D) Total vegetable score positively correlates with the abundance of *L. crispatus* and negatively correlates with the abundance of *L. iners* (E, Spearman Correlation).

### Higher Lactobacillus crispatus *dominance (CST-I) is associated with higher dietary quality and vegetable consumption*

We determine the vaginal CST for each participant. More than 40% exhibited CST-I (*L. crispatus* dominance), followed by CST-III (*L. iners* dominance), CST-II (*L. gasseri* dominance), CST-IV (non-*Lactobacillus*-dominated), and CST-V (*L. jensenii* dominance. **Table S4**) (27). We did not observe differences in CSTs by health status, IBD diagnosis (CD vs. UC), levels of fecal calprotectin, or any of the vaginal cytokines assessed (**Table S3 and S4**).

Pregnant individuals with CST-I (*L. crispatus*-dominated, N=21) showed a higher HEI-2015 score, reflective of better dietary quality than CST-III individuals (*L. iners*-dominated, N=14. **Figure 3)**. Moreover, scoring for vegetable and added sugar intakes was significantly higher in individuals with CST-I than those with CST-III (**Figure 3**).

## Discussion

### Principal findings

The vaginal and gut microbiota composition of pregnant individuals with IBD was comparable to HC during the third trimester of pregnancy. Yet pregnant individuals with IBD presented gut inflammation linked to high vaginal microbial diversity and a vaginal pro-inflammatory immune tone. Pregnant individuals with higher dietary quality and optimal consumption of vegetables exhibited a vaginal microbiota profile dominated by the beneficial *L. crispatus*.

### Results in the context of what is known

Reduced levels of IFN-γ and IL-4 and elevated levels of IL-8 and IL-6, as seen in the pregnant individuals with IBD in this study, have been associated with an increased risk of preterm birth (28). To the best of our knowledge, our study is the first one to assess vaginal inflammatory markers in IBD pregnant patients and might explain the increased risk of preterm birth of pregnant individuals with IBD, even when in remission or with mild disease (29, 30). Despite differences in the vaginal immune profile, the vaginal microbiota of pregnant individuals with IBD was comparable to healthy controls. This result is unexpected as inflammation relates to alteration in the microbiota in the vaginal mucosa (31, 32). Here, the vaginal immune profile was assessed by gene expression (by RT-qPCR) and no protein/cytokine levels; thus, further studies are required to accurately evaluate the immune tone in the vaginal mucosa.

We observe that gut inflammation in IBD individuals is positively associated with increased vaginal microbial diversity. Of note, contrary to the gut microbiota, high microbial diversity in the vagina relates to unhealthy states with increased risk for bacterial vaginosis (reviewed in (33)) and preterm birth (34, 35).

We then investigated whether diet impacts the vaginal microbiota. Pregnant individuals with higher dietary quality exhibited a vaginal microbiota profile dominated by the beneficial *L. crispatus,* whereas the vaginal microbiota profile of those with lower dietary quality was dominated by *L. iners*. For individual dietary components, our findings showed that high vegetable consumption was associated with greater microbial diversity (linked to vaginal dysbiosis (36)), similar to what previous studies have found for non-pregnant vegetarians compared to non-vegetarians (37). However, our results also showed that, despite the higher diversity, high vegetable intake resulted in a greater abundance of the beneficial *L. crispatus*. This indicates the importance of not only considering a diversity index but identifying members of the vaginal microbiome at the specie level to understand the potential implications of the microbiota in vaginal health.

Additionally, we found that lower added sugar intake resulted in decreased microbial alpha diversity and increased *L. crispatus*. Concomitantly, high vegetable consumption and low intake of sweetened beverages, has been also positively associated with abundance of *L. crispatus* in White and Black pregnant women (8).

Importantly, *L. crispatus* is believed to offer the most protective benefits to the host compared to other *Lactobacillus* species, with *L. iners* offering the least protective benefits (reviewed in (5)). *L. crispatus* creates a highly acidic vaginal niche (pH<4.5) inhospitable to non-beneficial microbes, such as bacterial vaginosis-related bacteria (38).

### Clinical implications

We find that intestinal inflammation correlates with high vaginal microbiota diversity, indicative of unhealthy states with increased risk for bacterial vaginosis and preterm birth. Thus, our results highlight the importance of continuing therapy during pregnancy to reduce IBD-related intestinal inflammation. Moreover, we found that diet can influence the dominance of a beneficial *L. crispatus* associated with decreased risk of pre-term birth and bacterial vaginosis. Hence, emphasizing dietary quality during pregnancy is a must, not only for the sustainment of pregnancy but to fuel a healthy vaginal microbiota.

### Research implications

Our results highlight the need for a large prospective study that includes pregnant individuals with IBD experiencing different disease activity (i.e., mild, moderate, severe). Future studies including dietary interventions will unveil the role of diet as a strategy to support a healthy vaginal microbiome.

### Strengths and limitations

Our study is constrained by a few key limitations. The modest sample size for both IBD and HC cohorts curtails the statistical comparisons, particularly when several confounding variables, such as antibiotic use, IBD medications, and gestational diabetes, are considered. This factor reduces the statistical power of the study. Additionally, the IBD samples predominantly represent individuals in remission or with mild disease with CD and only a few participants with UC, which narrows the scope of our conclusions to this specific severity level of IBD and IBD diagnosis. Moreover, the ethnic/racial composition of our study sample, which is mainly White, introduces a limitation since the vaginal microbiome is known to vary with race and ethnicity (39). These limitations suggest the need for studies that include participants with severe disease, an equal representation of IBD diagnosis (UC and CD), as well as individuals with diverse ethnic/racial backgrounds.

### Conclusions

Our results demonstrate that although the vaginal and gut microbiota of pregnant individuals with IBD and HC is similar in the third trimester of pregnancy, it varies depending on the immune tone of each mucosa. We show that pregnant individuals with IBD exhibit a pro-inflammatory cytokine profile that has been associated with an increased risk of pre-term birth. Finally, a high-quality diet with optimal intakes of vegetables and added sugars favors *L. crispatus* vaginal dominance.

## Acknowledgments

We thank all the participants for their time and effort in collecting samples and responding to surveys. We also thank Rafael Lopez Martinez for contributing to designing scripts for analysis automatization. This work was supported by the Leona M. and Harry B. Helmsley Charitable Trust.

## Data availability statement

The raw sequences and their associated metadata will be released at NCBI BioProject ID PRJNA915128 upon manuscript publication.

