## Supplemental for "Association of diet and inflammation with the vaginal microbiota of pregnant individuals with or without IBD"

**SUPPLEMENTARY MATERIAL**

**METHODS:**

**Recruitment:** We conducted a case–control study nested into our ongoing MELODY Trial, which is a prospective non-randomized diet intervention trial testing the effects of IBD anti-inflammatory diet during the third trimester of pregnancy on maternal IBD activity and microbiome composition in mothers and their babies(9). Written informed consent was obtained from all eligible participants. Eligible participants were adult (18+ years of age) pregnant individuals with or without IBD, with less than 29 weeks of pregnancy, carrying a singleton, with or without IBD diagnosis. We excluded pregnant individuals with scheduled C-section or induction of vaginal delivery prior to week 37 at the time of enrollment, those with medical conditions that required a special diet, or those unable to speak or understand English.

**Cytokine expression:** RNA from vaginal samples was reverse transcribed using iScript cDNA Synthesis Kit (Bio-Rad, USA), and qRT–PCRs were performed using iTaq Universal SYBR Green-Supermix (Bio-Rad, USA) in an Applied Biosystems ViiA7 Real-Time PCR machine (Thermo-Scientific, USA). Expression of each cytokine was measured in triplicate and the mean was normalized by the expression of a housekeeping gene (GAPDH). The $2^{(-\Delta\Delta Ct)}$ method was used to analyze the relative changes in gene expression.

**Dietary assessment:** We conducted three, 24-hour dietary recalls (24HDRs) around the same time of vaginal/stool collection. 24HDRs were performed using the USDA Automated Multiple Pass Method(40) in conjunction with the University of Minnesota Nutrition Data for Research (NDSR) software (Current version: NDS-R 2022)(41-44) as we have previously done(45-47). We estimated the Healthy Eating Index 2015 (HEI-2015) from the average of the three 24HDRs. The population ratio method(48) was used to compute the individual scores of each of the 13 HEI-2015 components: total fruits, whole fruits, total vegetables, greens and beans, dairy, total protein foods, seafood and plant proteins, fatty acids, refined grains, whole grains, sodium, added sugars and saturated fats. Six food groups have values from 0 to 5; and seven from 0 to 10, for a total maximum score of 100 as previously described(25). Higher values represent better dietary quality. Wilcoxon test was used to compare HEI-2015 and individual HEI-2015' dietary components by health status and CST.

31. Masson L, Barnabas S, Deese J, Lennard K, Dabee S, Gamieldien H, Jaumdally SZ, Williamson AL, Little F, Van Damme L, Ahmed K, Crucitti T, Abdellati S, Bekker LG, Gray G, Dietrich J, Jaspan H, Passmore JS.

2019. Inflammatory cytokine biomarkers of asymptomatic sexually transmitted infections and vaginal dysbiosis: a multicentre validation study. Sex Transm Infect 95:5-12.

32. Campisciano G, Zanotta N, Licastro D, De Seta F, Comar M. 2018. In vivo microbiome and associated immune markers: New insights into the pathogenesis of vaginal dysbiosis. Sci Rep 8:2307.

33. Chen X, Lu Y, Chen T, Li R. 2021. The female vaginal microbiome in health and bacterial vaginosis. Frontiers in cellular and infection microbiology:271.

34. Hillier SL, Nugent RP, Eschenbach DA, Krohn MA, Gibbs RS, Martin DH, Cotch MF, Edelman R, Pastorek JG, Rao AV. 1995. Association between bacterial vaginosis and preterm delivery of a low-birth-weight infant. New England Journal of Medicine 333:1737-1742.

35. Donders G, Van Calsteren K, Bellen G, Reybrouck R, Van den Bosch T, Riphagen I, Van Lierde S. 2009. Predictive value for preterm birth of abnormal vaginal flora, bacterial vaginosis and aerobic vaginitis during the first trimester of pregnancy. BJOG: An International Journal of Obstetrics & Gynaecology 116:1315-1324.

36. Srinivasan S, Hoffman NG, Morgan MT, Matsen FA, Fiedler TL, Hall RW, Ross FJ, McCoy CO, Bumgarner R, Marrazzo JM, Fredricks DN. 2012. Bacterial communities in women with bacterial vaginosis: high resolution phylogenetic analyses reveal relationships of microbiota to clinical criteria. PLoS One 7:e37818.

37. Song SD, Acharya KD, Zhu JE, Deveney CM, Walther-Antonio MRS, Tetel MJ, Chia N. 2020. Daily Vaginal Microbiota Fluctuations Associated with Natural Hormonal Cycle, Contraceptives, Diet, and Exercise. mSphere 5.

47. Zhao M, Chiriboga D, Olendzki B, Xie B, Li Y, McGonigal LJ, Maldonado-Contreras A, Ma Y. 2018. Substantial Increase in Compliance with Saturated Fatty Acid Intake Recommendations after One Year Following the American Heart Association Diet. Nutrients 10.

48. Kirkpatrick SI, Reedy J, Krebs-Smith SM, Pannucci TE, Subar AF, Wilson MM, Lerman JL, Tooze JA. 2018. Applications of the Healthy Eating Index for Surveillance, Epidemiology, and Intervention Research: Considerations and Caveats. J Acad Nutr Diet 118:1603-1621.

### SUPPLEMENTARY TABLES

**Table S1.** List of oligos used to determine vaginal cytokine gene expression.

| Serial No | Target gene (qRT primer) | Sequence (5' - 3') |
|---|---|---|
| 1 | IL-1β (F) | CAAGAGTGCTGGAGCGATAA |
| 2 | IL-1β (R) | CCGTTACAGCGGAGAGATATAAG |
| 3 | TNFα (F) | CACAGAAGACACTCAGGGAAAG |
| 4 | TNFα (R) | TCGAGGGAGTCACCCTTAAA |
| 5 | IL-6 (F) | GGAGGAGCCAAAGCTCAATAA |
| 6 | IL-6 (R) | ACTCCACACCACAAGAAGATG |
| 7 | IL-4 (F) | CCTGCTTCATGAGGGAAACT |
| 8 | IL-4 (R) | GGTGACAGAACAAGACCCTATC |
| 9 | IL-8 (F) | GAAAGGAAGTAGCTGGCAGAG |
| 10 | IL-8 (R) | GGGTGGAAAGGTTTGGAGTAT |
| 11 | IL-10 (F) | TGGAGTGAGTCCTGGAGAAATA |
| 12 | IL-10 (R) | CTCCATGTCCATCACACTTAGG |
| 13 | IFN-γ (F) | CCAGCCATCCTCAGAAATGT |
| 14 | IFN-γ (R) | CTTGCATCTCCTCACTCTAACC |
| 15 | IL-17 (F) | CCATAGTGAAGGCAGGAATCA |
| 16 | IL-17 (R) | GAGGTGGATCGGTTGTAGTAATC |
| 17 | GM-CSF (F) | CCTCCAGCAGGAATGTCTTAAT |
| 18 | GM-CSF (R) | TCTGACTCCTTGGGATGAAATG |
| 19 | GAPDH (F) | GTGCTCCCACTCCTGATTT |
| 20 | GAPDH (R) | CCTTCTCTAAGTCCCTCCTACA |

**Table S2.** Overview number of sequences and amplicon sequence variants (ASVs) obtained from the vaginal samples and stool samples.

| Parameter | Vaginal samples | | | Stool samples | |
|---|---|---|---|---|---|
| | **Raw sequences** | **Post-filtering** | **Post-filtering/rarefaction*** | **Raw sequences** | **Rarefied sequences**** |
| N samples | 48 | 48 | 48 | 45 | 44 |
| Total number of seqs | 2,751,605 | 2,742,919 | 336,000 | 652,433,458 | 236,170,044 |
| Mean seqs/sample | 57,325 | 57,144 | 7,000 | 14,498,521 | 5,367,501 |
| Standard deviation of seqs/sample | 24,010 | 23,945 | - | 7,468,723 | - |
| Max number of seqs in a sample | 123,825 | 123,720 | 7,000 | 30,294,951 | 5,367,501 |
| Min number of seqs in a sample | 12,916 | 12,872 | 7,000 | 715 | 5,367,501 |
| Total number of different ASVs or Strains | 1,425 | 573 | 563 | 1,420 | 897 |
| Mean ASV or Strains/sample | 48 | 36 | 33 | 115 | 118 |
| Standard deviation of ASV or Strains/sample | 38 | 23 | 23 | 46 | 43 |
| Max number of ASVs or Strains in a sample | 208 | 109 | 105 | 249 | 249 |
| Min number of ASVs or Strains in a sample | 6 | 6 | 5 | 1 | 57 |

* Sequences were rarefied at *7,000 sequences/sample or **5,367,501 sequences/sample, each representing the highest number of sequences that included all vaginal and stool samples, respectively.

**Table S3.** Demographic and clinical variables for pregnant individuals with Crohn's Disease (CD) or Ulcerative Colitis (UC) recruited for this study between 2019 and 2022.

| **Demographics and clinical variables** | **CD (N=18)** | **UC (N=5)** | **Overall (N=23)** | **P value[&]** |
|---|---|---|---|---|
| **Age** | | | | *0.822* |
| Mean (SD) | 33.2 (4.86) | 33.4 (4.22) | 33.3 (4.63) | |
| Median [Min, Max] | 33.5 [22.0, 41.0] | 32.0 [28.0, 39.0] | 33.0 [22.0, 41.0] | |
| **BMI categories [&&]** | | | | *0.126* |
| Underweight | 1 (5.6%) | 0 (0%) | 1 (4.3%) | |
| Normal | 10 (55.6%) | 2 (40.0%) | 12 (52.2%) | |
| Overweight | 6 (33.3%) | 2 (40.0%) | 8 (34.8%) | |
| Obese | 1 (5.6%) | 1 (20.0%) | 2 (8.7%) | |
| **Race** | | | | - |
| White | 18 (100%) | 5 (100%) | 23 (100%) | |
| Asian | 0 (0%) | 0 (0%) | 0 (0%) | |
| Other | 0 (0%) | 0 (0%) | 0 (0%) | |
| **Gestational diabetes** | | | | *0.053* |
| Yes | 1 (5.6%) | 2 (40.0%) | 3 (13.0%) | |
| No | 14 (77.8%) | 2 (40.0%) | 16 (69.6%) | |
| Information unavailable | 3 (16.7%) | 1 (20.0%) | 4 (17.4%) | |
| **Use of antibiotic during pregnancy** | | | | *1.000* |
| No | 15 (83.3%) | 4 (80.0%) | 19 (82.6%) | |
| Yes | 3 (16.7%) | 0 (0%) | 3 (13.0%) | |
| Information unavailable | 0 (0%) | 1 (20.0%) | 1 (4.3%) | |
| **Disease activity [&&&]** | | | | *0.456* |
| Mild disease | 5 (27.8%) | 1 (20.0%) | 6 (26.1%) | |
| Remission | 10 (55.6%) | 3 (60.0%) | 13 (56.5%) | |
| Information unavailable | 3 (16.7%) | 1 (20.0%) | 4 (17.4%) | |
| **Use of IBD medication** | | | | *0.342* |
| No | 6 (33.3%) | 3 (60.0%) | 9 (39.1%) | |
| Yes | 12 (66.7%) | 2 (40.0%) | 14 (60.9%) | |
| **Preterm** | | | | *0.272* |
| No | 12 (66.7%) | 3 (60.0%) | 15 (65.2%) | |
| Yes | 2 (11.1%) | 2 (40.0%) | 4 (17.4%) | |
| Missing | 4 (22.2%) | 0 (0%) | 4 (17.4%) | |
| **Infant birth weight (g)** | | | | *0.411* |
| Mean (SD) | 3100 (515) | 3330 (169) | 3150 (470) | |
| Median [Min, Max] | 3230 [1810, 3710] | 3230 [3180, 3540] | 3230 [1810, 3710] | |
| **Fecal calprotectin (ng/mg)** | | | | *1.000* |
| Mean (SD) | 69.4 (65.9) | 59.5 (42.7) | 67.3 (60.9) | |
| Median [Min, Max] | 45.2 [3.11, 208] | 45.2 [6.89, 111] | 45.2 [3.11, 208] | |
| **Community State Types (CSTs)** | | | | *0.209* |
| I | 7 (38.9%) | 2 (40.0%) | 9 (39.1%) | |
| II | 4 (22.2%) | 0 (0%) | 4 (17.4%) | |
| III | 2 (11.1%) | 3 (60.0%) | 5 (21.7%) | |
| IV-C | 3 (16.7%) | 0 (0%) | 3 (13.0%) | |

| | | | |
|---|---|---|---|
| V | 2 (11.1%) | 0 (0%) | 2 (8.7%) |

&Fisher's exact test for categorical variables and Wilcoxon test for continuous variables.

&&BMIs categories correspond to the WHO’s classifications: Underweight (<18.5), normal weight (18.5–24.9), overweight (≥25.0), and obese (≥30).

&&&Disease activity was estimated using the Harvey Bradshaw Index and the Mayo score for individuals with Crohn's Disease or Ulcerative colitis, respectively.

**Table S4.** Vaginal microbial Community State Types (CSTs) distribution by health status, levels of fecal calprotectin, and expression of vaginal cytokines

| Variables | CST-I (N=21) | CST-II (N=7) | CST-III (N=14) | CST-IV-C (N=4) | CST-V (N=2) | Overall (N=48) | P value& |
|---|---|---|---|---|---|---|---|
| **Health status** | | | | | | | *0.369* |
| HC | 12 (57.1%) | 3 (42.9%) | 9 (64.3%) | 1 (25.0%) | 0 (0%) | 25 (52.1%) | |
| IBD | 9 (42.9%) | 4 (57.1%) | 5 (35.7%) | 3 (75.0%) | 2 (100%) | 23 (47.9%) | |
| **Fecal calprotectin (ng/mg)** | | | | | | | *0.373* |
| Mean (SD) | 37.0 (55.8) | 36.6 (47.2) | 26.3 (26.5) | 40.4 (47.8) | 120 (124) | 37.6 (51.1) | |
| Median | 8.54 | 12.4 | 19.4 | 19.5 | 120 | 15.3 | |
| **IL-6** | | | | | | | *0.114* |
| Mean (SD) | 3.8 (4.1) | 4.4 (3.4) | 2.9 (2.3) | 7.5 (3.9) | 8.7 (0.05) | 4.1 (3.7) | |
| Median | 2.14 | 4.05 | 2.5 | 7.2 | 8.76 | 3.12 | |
| **IL-4** | | | | | | | *0.474* |
| Mean (SD) | 0.761 (0.658) | 0.850 (0.762) | 0.855 (0.684) | 0.377 (0.301) | 0.193 (0.140) | 0.746 (0.651) | |
| Median | 0.421 | 0.877 | 0.581 | 0.277 | 0.193 | 0.428 | |
| **IL-1** | | | | | | | *0.086* |
| Mean (SD) | 1.22 (0.453) | 1.18 (0.403) | 1.04 (0.392) | 0.763 (0.519) | 2.01 (0.321) | 1.16 (0.469) | |
| Median | 1.23 | 1.06 | 1.02 | 0.655 | 2.01 | 1.06 | |
| **TNF-alpha** | | | | | | | *0.237* |
| Mean (SD) | 3.46 (3.14) | 4.42 (4.05) | 3.00 (2.51) | 2.48 (1.90) | 11.2 (1.79) | 3.71 (3.33) | |
| Median | 1.32 | 4.74 | 1.69 | 1.93 | 11.2 | 2.1 | |
| **IFN-gamma** | | | | | | | *0.431* |
| Mean (SD) | 0.583 (0.671) | 0.663 (0.951) | 0.776 (0.751) | 0.152 (0.303) | 0.00340 (0.00157) | 0.591 (0.715) | |
| Median | 0.561 | 0.001 | 0.712 | 0.000763 | 0.0034 | 0.511 | |
| **IL-8** | | | | | | | *0.201* |
| Mean (SD) | 3.05 (3.05) | 3.96 (4.26) | 2.25 (2.12) | 4.15 (4.24) | 11.5 (2.62) | 3.39 (3.48) | |
| Median | 1.57 | 1.26 | 1.79 | 2.68 | 11.5 | 1.93 | |
| **GM-CSF** | | | | | | | *0.076* |
| Mean (SD) | 1.04 (0.453) | 1.10 (0.561) | 1.15 (0.488) | 0.689 (0.565) | 4.31 (0.303) | 1.19 (0.815) | |
| Median | 1.02 | 0.917 | 1.08 | 0.485 | 4.31 | 1.02 | |
| **IL-10** | | | | | | | *0.176* |
| Mean (SD) | 1.15 (0.625) | 1.27 (0.745) | 1.20 (0.496) | 1.09 (0.500) | 5.59 (0.473) | 1.36 (1.06) | |
| Median | 1 | 1.17 | 1.11 | 0.922 | 5.59 | 1.09 | |
| **IL-17** | | | | | | | *0.096* |
| Mean (SD) | 1.93 (1.20) | 2.60 (1.31) | 1.59 (1.08) | 1.85 (1.26) | 6.03 (1.09) | 2.09 (1.44) | |
| Median | 2.07 | 2.49 | 1.17 | 2.08 | 6.03 | 1.99 | |

**Table S5.** HEI-2015 score and individual dietary components scoring distribution by health status.

| HEI-2015 dietary components (scoring scale) | IBD (N=22) | HC (N=25) | Overall (N=47) | P value& |
|---|---|---|---|---|
| **Healthy Eating Index (0-100)** | | | | *0.175* |
| Mean (SD) | 66.6 (13.8) | 61.4 (13.2) | 63.8 (13.6) | |
| Median [Min, Max] | 67.5 [45.0, 91.0] | 64.0 [35.0, 84.0] | 65.0 [35.0, 91.0] | |
| **Total vegetable (0-5)** | | | | *0.361* |
| Mean (SD) | 3.68 (1.39) | 4.20 (1.12) | 3.96 (1.27) | |
| Median [Min, Max] | 4.00 [1.00, 5.00] | 5.00 [1.00, 5.00] | 5.00 [1.00, 5.00] | |
| **Greens and beans (0-5)** | | | | *0.179* |
| Mean (SD) | 4.41 (1.47) | 3.48 (1.90) | 3.91 (1.75) | |
| Median [Min, Max] | 5.00 [0, 5.00] | 5.00 [0, 5.00] | 5.00 [0, 5.00] | |
| **Total fruit (0-5)** | | | | *0.279* |
| Mean (SD) | 3.23 (1.57) | 2.80 (1.58) | 3.00 (1.57) | |
| Median [Min, Max] | 3.50 [0, 5.00] | 3.00 [0, 5.00] | 3.00 [0, 5.00] | |
| **Whole fruit (0-5)** | | | | *0.281* |
| Mean (SD) | 3.82 (1.76) | 3.44 (1.87) | 3.62 (1.81) | |
| Median [Min, Max] | 5.00 [0, 5.00] | 5.00 [0, 5.00] | 5.00 [0, 5.00] | |
| **Whole grains (0-10)** | | | | *0.728* |
| Mean (SD) | 5.36 (3.27) | 5.52 (3.27) | 5.45 (3.24) | |
| Median [Min, Max] | 5.50 [0, 10.0] | 6.00 [0, 10.0] | 6.00 [0, 10.0] | |
| **Dairy (0-10)** | | | | *0.659* |
| Mean (SD) | 6.68 (2.23) | 6.76 (2.60) | 6.72 (2.41) | |
| Median [Min, Max] | 6.00 [3.00, 10.0] | 7.00 [3.00, 10.0] | 6.00 [3.00, 10.0] | |
| **Total protein foods (0-5)** | | | | *0.508* |
| Mean (SD) | 4.64 (0.727) | 4.32 (1.03) | 4.47 (0.905) | |
| Median [Min, Max] | 5.00 [3.00, 5.00] | 5.00 [1.00, 5.00] | 5.00 [1.00, 5.00] | |
| **Seafood and plant protein (0-5)** | | | | *0.212* |
| Mean (SD) | 3.86 (1.83) | 3.36 (1.78) | 3.60 (1.80) | |
| Median [Min, Max] | 5.00 [0, 5.00] | 3.00 [0, 5.00] | 5.00 [0, 5.00] | |
| **Fatty acid ratio (0-10)** | | | | *0.162* |
| Mean (SD) | 5.50 (3.60) | 3.04 (2.94) | 4.19 (3.46) | |
| Median [Min, Max] | 5.00 [0, 10.0] | 2.00 [0, 10.0] | 3.00 [0, 10.0] | |
| **Sodium (0-10)** | | | | *0.217* |
| Mean (SD) | 5.84 (2.79) | 5.82 (3.22) | 5.83 (2.97) | |
| Median [Min, Max] | 6.00 [1.00, 10.0] | 6.50 [0, 10.0] | 6.00 [0, 10.0] | |
| **Refined grains (0-10)** | | | | *0.251* |
| Mean (SD) | 6.09 (4.10) | 6.76 (2.79) | 6.45 (3.44) | |
| Median [Min, Max] | 8.00 [0, 10.0] | 7.00 [0, 10.0] | 8.00 [0, 10.0] | |
| **Added sugar (0-10)** | | | | *0.514* |
| Mean (SD) | 8.18 (1.79) | 8.20 (1.91) | 8.19 (1.84) | |
| Median [Min, Max] | 9.00 [5.00, 10.0] | 9.00 [5.00, 10.0] | 9.00 [5.00, 10.0] | |
| **Saturated fats (0-10)** | | | | *0.847* |
| Mean (SD) | 5.23 (3.32) | 3.48 (3.00) | 4.30 (3.24) | |
| Median [Min, Max] | 5.50 [0, 10.0] | 3.00 [0, 10.0] | 4.00 [0, 10.0] | |

& Fisher's exact test for categorical variables and Wilcoxon test for continuous variables

**Table S6.** HEI-2015 score and individual dietary components scoring distribution by Community State Types (CSTs).

| Diet variables | Community State Types<br>I<br>(N=21) | <br>II<br>(N=6) | <br>III<br>(N=14) | <br>IV-C<br>(N=4) | <br>V<br>(N=2) | Overall<br>(N=47) | P value[&] |
|---|---|---|---|---|---|---|---|
| *Healthy Eating Index (total score 0-100)* | | | | | | | *0.098* |
| Mean (SD) | **68.4 (11.3)** | 61.0 (15.7) | **55.2 (13.7)** | 69.0 (9.49) | 73.5 (13.4) | 63.8 (13.6) | |
| Median | **67** | 65.5 | **53** | 65.5 | 73.5 | 65 | |
| *Total vegetables (0-5)* | | | | | | | *0.007* |
| Mean (SD) | **4.57 (0.978)** | 3.50 (1.38) | **3.21 (1.12)** | 4.25 (1.50) | 3.50 (2.12) | 3.96 (1.27) | |
| Median | **5** | 4 | **3** | 5 | 3.5 | 5 | |
| *Greens and Beans (0-5)* | | | | | | | *0.169* |
| Mean (SD) | 4.38 (1.36) | 3.50 (2.07) | 3.07 (2.20) | 4.75 (0.500) | 4.50 (0.707) | 3.91 (1.75) | |
| Median | 5 | 4.5 | 4 | 5 | 4.5 | 5 | |
| *Total fruit (0-5)* | | | | | | | *0.016* |
| Mean (SD) | 2.86 (1.31) | 3.33 (1.86) | 2.29 (1.59) | 4.75 (0.500) | 5.00 (0) | 3.00 (1.57) | |
| Median | 3 | 3.5 | 2 | 5 | 5 | 3 | |
| *Whole fruit (0-5)* | | | | | | | *0.271* |
| Mean (SD) | 3.81 (1.72) | 3.83 (1.60) | 2.79 (2.12) | 4.50 (1.00) | 5.00 (0) | 3.62 (1.81) | |
| Median | 5 | 4.5 | 3 | 5 | 5 | 5 | |
| *Whole grains (0-10)* | | | | | | | *0.947* |
| Mean (SD) | 5.52 (3.33) | 5.33 (3.14) | 5.14 (3.35) | 5.50 (4.43) | 7.00 (0) | 5.45 (3.24) | |
| Median | 6 | 5.5 | 4.5 | 6 | 7 | 6 | |
| *Dairy (0-10)* | | | | | | | *0.919* |
| Mean (SD) | 6.62 (2.52) | 6.00 (2.45) | 6.93 (2.53) | 7.25 (1.71) | 7.50 (3.54) | 6.72 (2.41) | |
| Median | 6 | 6 | 6.5 | 7.5 | 7.5 | 6 | |
| *Total protein foods (0-5)* | | | | | | | *0.482* |
| Mean (SD) | 4.33 (1.11) | 5.00 (0) | 4.50 (0.760) | 4.50 (0.577) | 4.00 (1.41) | 4.47 (0.905) | |
| Median | 5 | 5 | 5 | 4.5 | 4 | 5 | |
| *Seafood and plant protein (0-5)* | | | | | | | *0.112* |
| Mean (SD) | 4.24 (1.37) | 3.67 (2.16) | 2.79 (1.89) | 2.50 (2.38) | 4.50 (0.707) | 3.60 (1.80) | |
| Median | 5 | 5 | 2 | 2.5 | 4.5 | 5 | |
| *Fatty acid ratio (0-10)* | | | | | | | *0.306* |
| Mean (SD) | 4.95 (3.34) | 2.67 (3.27) | 3.36 (3.52) | 5.25 (3.20) | 4.50 (6.36) | 4.19 (3.46) | |
| Median | 4 | 1.5 | 2 | 4 | 4.5 | 3 | |
| *Sodium (0-10)* | | | | | | | *0.305* |
| Mean (SD) | 6.57 (2.60) | 6.00 (3.35) | 4.57 (3.27) | 5.00 (2.94) | 8.00 (1.41) | 5.83 (2.97) | |
| Median | 7 | 7 | 3.5 | 5.5 | 8 | 6 | |
| *Refined grains (0-10)* | | | | | | | *0.627* |
| Mean (SD) | 6.62 (3.80) | 6.83 (2.86) | 5.36 (3.50) | 7.50 (2.52) | 9.00 (1.41) | 6.45 (3.44) | |
| Median | 8 | 6.5 | 5 | 8 | 9 | 8 | |
| *Added sugar (0-10)* | | | | | | | *0.02* |
| Mean (SD) | **9.10 (1.51)** | 8.00 (1.79) | **7.07 (1.82)** | 7.75 (2.06) | 8.00 (1.41) | 8.19 (1.84) | |
| Median | **10** | 8.5 | **6.5** | 7.5 | 8 | 9 | |

| | | | | | | | |
|---|---|---|---|---|---|---|---|
| *Saturated fats (0-10)* | | | | | | | *0.532* |
| Mean (SD) | 5.10 (3.46) | 2.67 (2.50) | 3.79 (3.04) | 4.75 (3.30) | 3.50 (4.95) | 4.30 (3.24) | |
| Median | 5 | 2.5 | 3.5 | 6 | 3.5 | 4 | |

& Kruskal-Wallis test followed by Wilcoxon tests for significant variables; in bold are indicated the pairwise analyses yielding significant differences between groups.

### SUPPLEMENTARY FIGURES

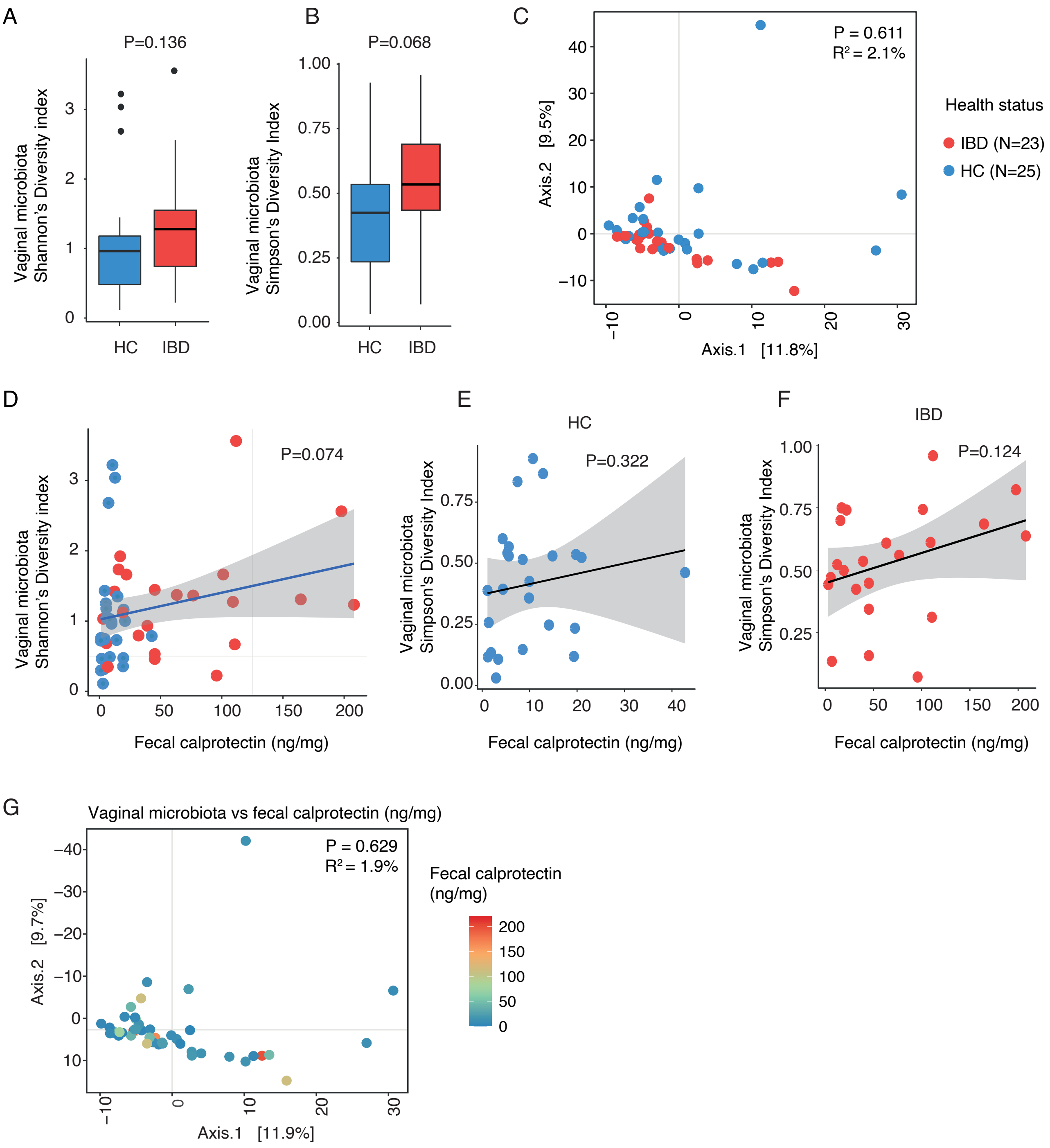


**Figure S1.** Comparison of vaginal microbiota diversity by health status and fecal calprotectin levels. Shannon (A) and Simpson's (B) diversity indices to evaluate the microbial diversity. A linear regression model assessed the comparison by health status. A Principal Coordinates Analysis (PCoA) (C) was performed based on the Aitchison distance. The Shannon diversity index (D) by health status, with linear regression modeling for analysis. Simpson's diversity index was employed separately for Healthy Controls

(E) and IBD patients (F), indicating no significant correlations; linear regression models were utilized for these assessments. For the PCoA, PERMANOVA was applied to calculate P values and $R^2$, helping quantify the explained variance.

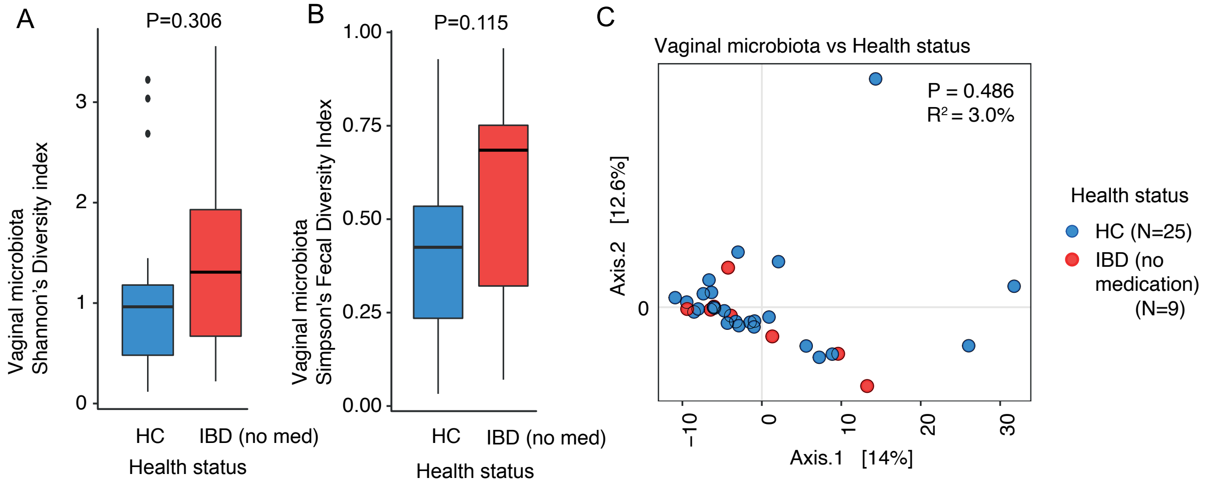


**Figure S2**. Comparison of vaginal microbial diversity in IBD patients off medication vs Healthy Controls (HC). Shannon's index (A) and Simpson's index (B). The significance levels were computed using a linear regression model. Beta diversity patterns are depicted through Principal Coordinates Analysis (PCoA) based on Aitchison distances (C), statistical significance and explained variance are reported through P values and $R^2$, using PERMANOVA.

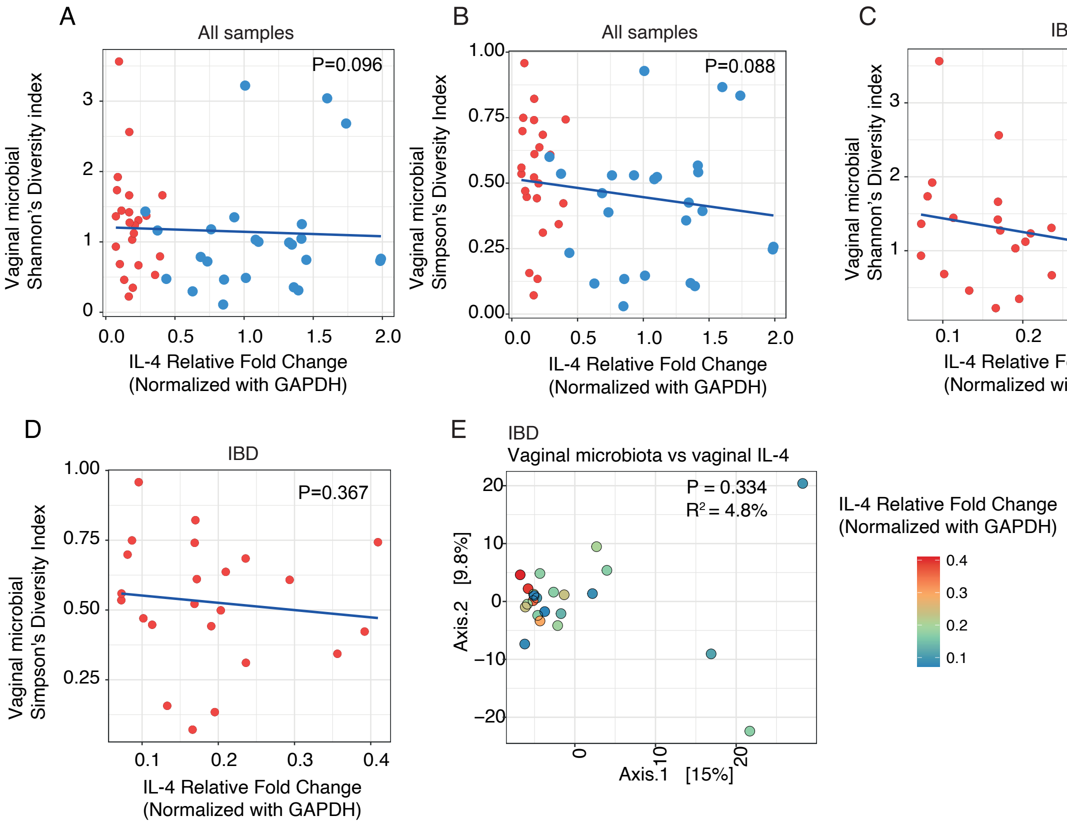


**Figure S3**. Vaginal microbial diversity and its of association with vaginal IL-4. Vaginal microbial diversity using Shannon (A) and Simpson's (B) diversity index by vaginal IL-4 relative fold change among all samples, reveals no significant associations, with linear regression models. Same analysis, although only among IBD patients using Shannon (C) and Simpson's (D) diversity index by vaginal IL-4 relative fold change, was not significant either. Principal Coordinates Analysis (PCoA), based on Aitchison distance, demonstrates no significance for vaginal microbial composition by IL-4 including only IBD patients (E). For PCoA PERMANOVA was used to obtain the P valued and to quantify the variance explained, as shown by P values and $R^2$.

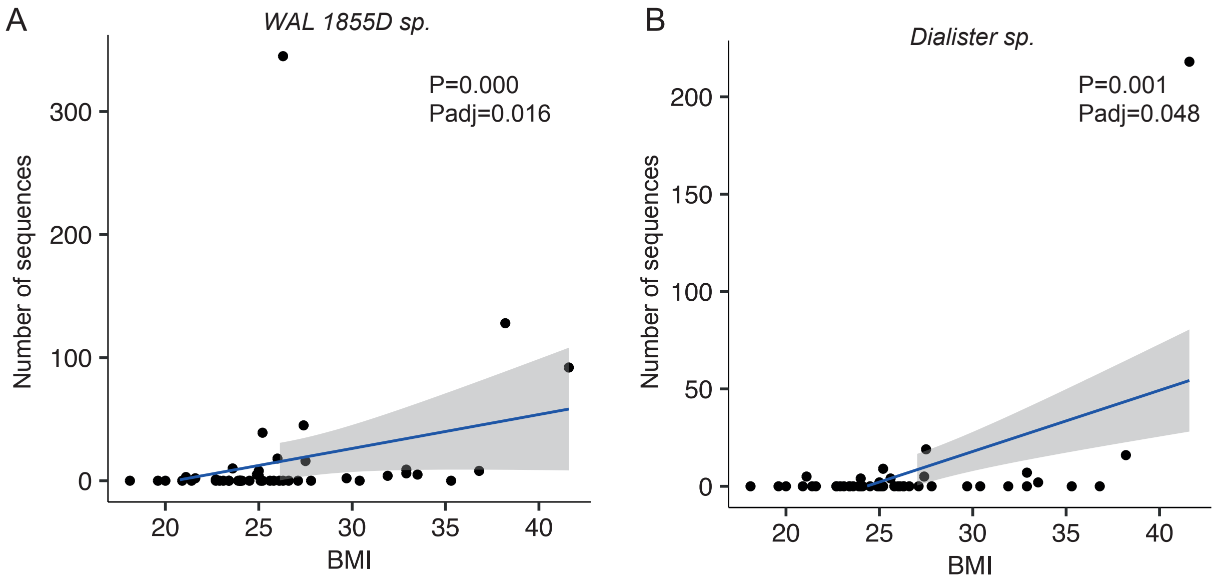


**Figure S4.** Vaginal species positively correlated with women's BMI. Number sequences by BMI for WAL 1855D sp (A) and *Dialister* sp (B). Analysis was performed with MaAsLin2 algorithm. P values were adjusted for multiple comparisons.

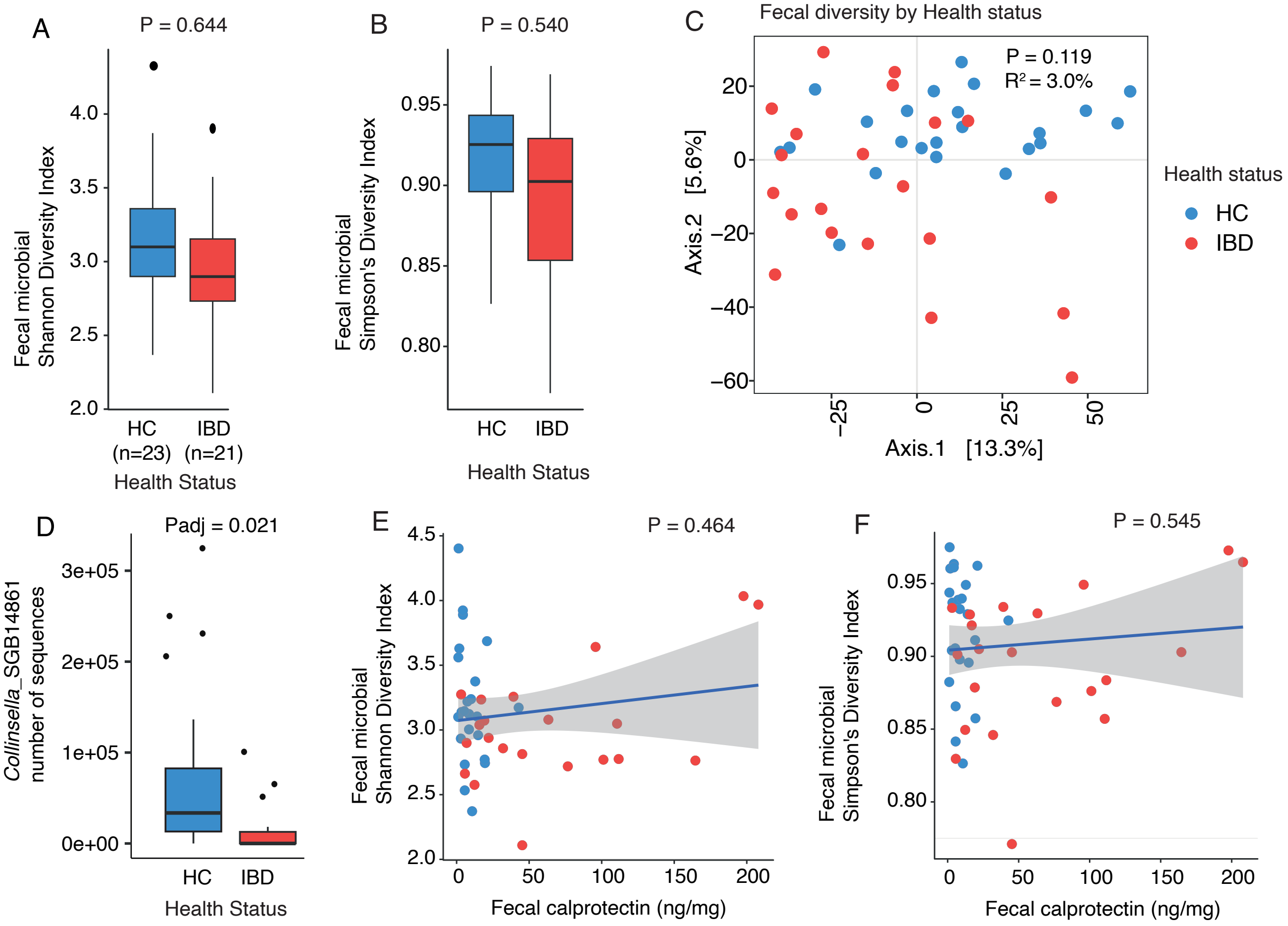


**Figure S5.** Comparison of fecal microbiota diversity by health status and fecal calprotectin levels. Shannon (A) and Simpson's (B) diversity indices to evaluate the microbial diversity. A linear regression model assessed the comparison by health status. A Principal Coordinates Analysis (PCoA) was performed based on the Aitchison distance to compare health status (C). Taxa significantly different by health status based on the MaAsLin2 algorithm (D). Shannon (E) and Simpson's (F) diversity indexes by fecal calprotectin levels, with linear regression modeling for analysis. For the PCoA, PERMANOVA was applied to calculate P values and $R^2$, helping quantify the explained variance

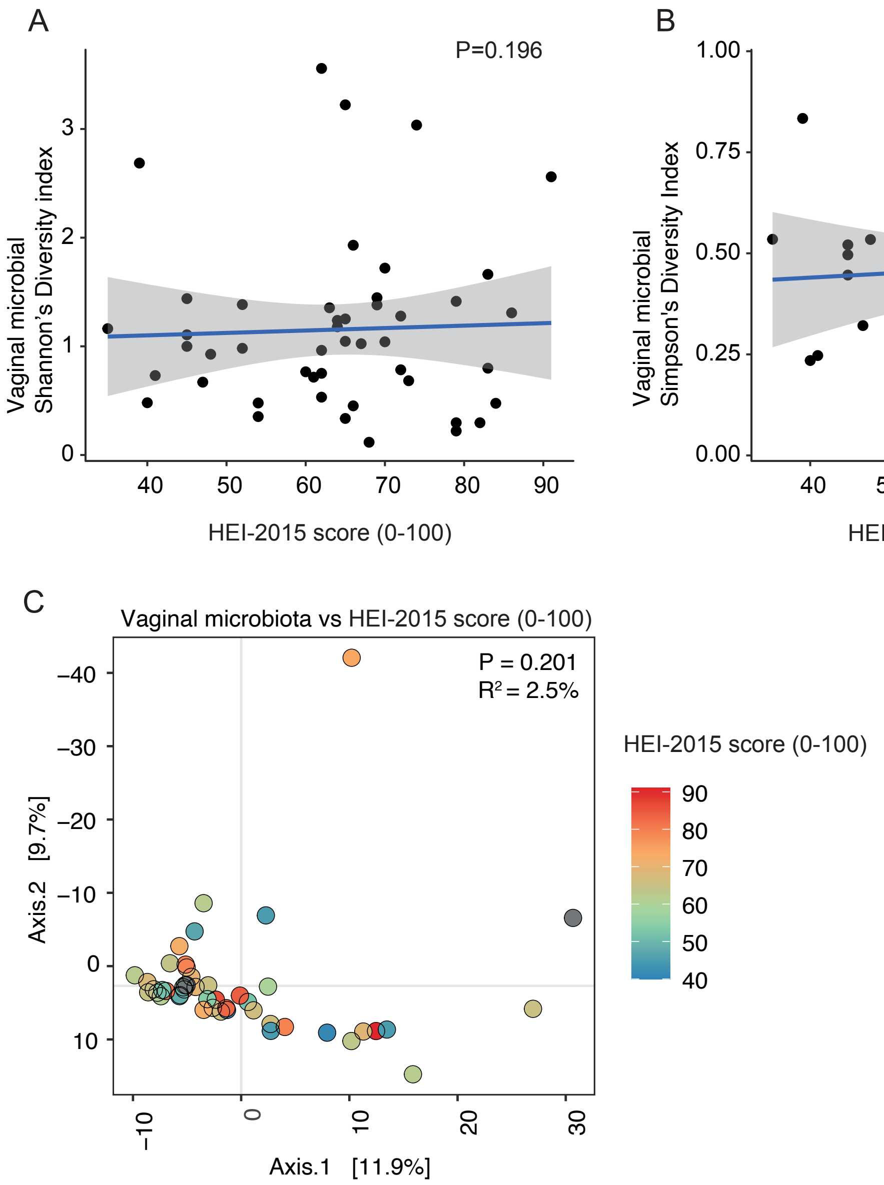


**Figure S6.** Diet quality is not associated with vaginal microbial alpha or beta diversity. HEI-2015 score for alpha diversity measured with Shannon's (A) and Simpson's (B) diversity indexes. P values were calculated with a linear regression model. Principal Coordinates Analysis (PCoA), based on Aitchison distance, for vaginal microbial composition by HEI-2015 score (B). PERMANOVA was used to obtain the P valued and to quantify the variance explained, as shown by P values and R2.

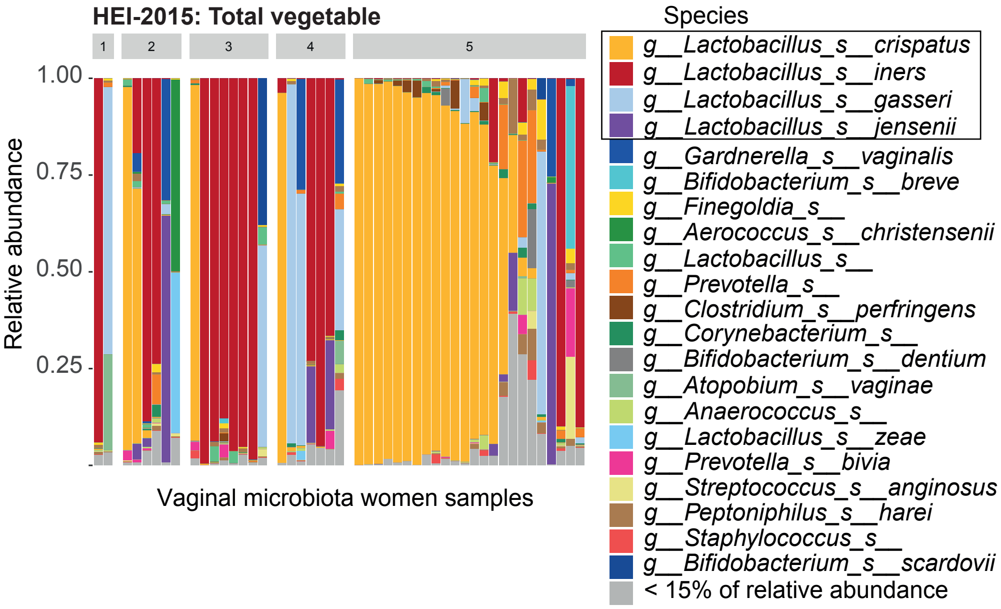


**Figure S7.** Microbial profile of pregnant individuals included in the study grouped by HEI-2015 total vegetable score and sorted by *L. crispatus* relative abundance. Each bar represents a unique sample.
